# Dual Specificity Phosphatase 3 knockdown drives myeloid leukemia cells to differentiate into macrophages and polarize

**DOI:** 10.1101/2024.09.30.615030

**Authors:** Jessica O. Farias, Diana R.D.C.G. Pacheco, Yuli T. Magalhaes, Lilian C. Russo, Viktor K. Boell, Donna J.F. Hilares, Fabio L. Forti

**Affiliations:** Laboratory of Signaling in Biomolecular Systems, Department of Biochemistry, Institute of Chemistry, University of São Paulo, Sao Paulo-SP, Brazil

**Keywords:** Protein Tyrosine Phosphatases, DUSP3, Nucleophosmin, Myeloid leukemia cells, Cell differentiation, Macrophages polarization

## Abstract

The dual-specificity phosphatase 3 (DUSP3) has been implicated in the maintenance of genomic stability, cell cycle, proliferation, and differentiation. Recently we reported an important role of the interaction between DUSP3 and nucleophosmin (NPM) proteins on the regulation of the p53 actions to maintain genomic stability. Since both p53 and NPM often have mutations related to a diverse set of leukemia, this work aimed to evaluate the roles of DUSP3 in the differentiation of two acute myeloid leukemia cell lines not expressing the p53 protein, and the potential correlations with NPM expression. The results demonstrated higher levels of DUSP3 in THP-1 cells compared to HL-60 cells under basal conditions. After PMA-induced differentiation into macrophages, only HL-60 cells presented a dramatic decrease in DUSP3 and NPM proteins expression. The permanent DUSP3 knockdown in THP-1 and HL-60 cells contributed to their differentiation and non-classical polarization after PMA exposure, since the CD14, MHCII, and CD163 markers were decreased whereas the CD11b and CD206 markers were increased. Bioinformatics analyses identified that the negative regulation of the *npm1* and *dusp3* genes correlates with the reduced survival of patients with acute myeloid leukemia (AML) and the strong positive correlation existing between the expression of these two genes is progressively lost according to the degree of maturation of the myeloid cells. These results suggest DUSP3 plays regulatory roles of differentiation and polarization of myeloid cells, and its association with NPM expression levels may allow a better understanding of mechanisms involved in leukemia and treatment resistance.

**HIGHLIGHTS:** DUSP3 knockdown drives myeloid leukemia cells to differentiation

DUSP3 silencing drives myeloid leukemia cells to macrophage polarization

DUSP3 and NPM association are potential targets for leukemia treatment and resistance

## 1. INTRODUCTION

Acute myeloid leukemia (AML) is a heterogeneous and aggressive hematopoietic neoplasm characterized by the infiltration of bone marrow, blood and other tissues by highly proliferative cells (undifferentiated precursors), resulting in impaired hematopoiesis and bone marrow failure ^1^. Mutations in the nucleophosmin 1 (*npm1*) gene are the most frequent in AML, occurring in 25-30% of patients, leading to aberrant expression of the NPM protein, which is retained in the cytoplasm, stimulating proliferation and the development of leukemia^1^. NPM is a nucleolar/nuclear phosphoprotein that regulates several distinct processes such as ribosomal biogenesis ^2^, centrosome duplication ^3^, and the maintenance of genomic integrity through the control of chromosomal ploidy, in addition to playing an important indirect participation in DNA repair ^2,4,5^. In this scenario, our group demonstrated for the first time that dual-specificity phosphatase 3 (DUSP3) interacts with NPM and other nuclear proteins such as nucleolin, nibrin, hnRNP C1/C2, and ATM/ATR in conditions of genotoxic stress ^6,7^. Additionally, we recently proved that DUSP3 can modulate the translation of IRES-dependent mRNAs by dephosphorylating hnRNP C1/C2, which affects the stability and interaction of protein complexes with RNA recognition motifs (RRM) and RNA following genotoxic stress ^8^.

DUSP3 is a class I protein tyrosine phosphatase belonging to the atypical dual phosphatases group ^8^, overexpressed in many types of cancer, such as breast ^9^ and prostate cancer^10^, while its mRNA and protein levels were markedly underexpressed in corresponding adjacent healthy tissues, as wells as in patients with non-small cell lung cancer (NSCLC) ^11^. The DUSP3 expression was highly detected in macrophages, as well as in immature dendritic cells, mast cells and neutrophils of mice under basal conditions, according to the Reference Genomics Database of Immune Cell (RefDIC) ^12^. Over the past years, protein phosphatases have gained relevance in the differentiation of leukemic cells, such as the serine/threonine phosphatase PPM1A ^13^, and the dual specificity tyrosine phosphatases Pyst2 ^14^ and DUSP5 ^15^. Some studies have reported roles of DUSP3 in cells of the immune system, particularly in the T cell signaling pathway ^16,17^, while others suggested a role for DUSP3 in cell differentiation due to its action upon MAPKs ^18^, although this has not yet been extensively documented in leukemia models.

Therefore, considering the roles of NPM in leukemic cells and our previous findings of the DUSP3-NPM interaction, we investigated the influence of DUSP3 knockdown and NPM expression levels on the differentiation and activation of two models of acute myeloid leukemia cells, THP-1 and HL-60 cell lines, which represent different stages of maturation, both with or without macrophage-induced differentiation stimuli by phorbol 12-myristate 13-acetate (PMA). Moreover, through a bioinformatics analysis, we examined the impact of *dusp3* and *npm1* gene expression on the survival of patients with AML, as well as the potential correlations between these genes and others known to be involved in myeloid cell differentiation.

## 2. Materials & Methods

### 2.1. Cell culture

HL-60 and THP-1 cell lines were maintained in RPMI (Roswell Park Memorial Institute) medium supplemented with 10% fetal bovine serum (Cultilab), 0.025 g/L penicillin (Sigma-Aldrich), and 0.1 g/L streptomycin (Life Technologies) at 37°C in humidified atmosphere of 5% CO_2_ and 95% air. For the THP-1 cells, the medium was supplemented with 0.05 mM β-mercaptoethanol.

### 2.2. Cell differentiation

THP-1 and HL-60 cells were differentiated into macrophages using phorbol 12-myristate 13-acetate (PMA) (Sigma-Aldrich) at concentrations of 5 ng/mL and 100 ng/mL, respectively, for 48 hours. CD14 expression served as the differentiation control, assessed before and after PMA treatments.

### 2.3. Western blotting

To obtain total cell lysates, cells were centrifuged at 800 rpm for 5 minutes at 4°C, washed twice with PBS, and centrifuged in the same conditions. The cell pellet was resuspended in RIPA buffer (50 mM Tris-HCl pH 7.2, 1% Triton X-100, 0.5% sodium deoxycholate, 0.1% SDS, 500 mM NaCl, 10 mM MgCl_2_) containing protease inhibitors (2 μg/mL leupeptin, pepstatin A and aprotinin, 1 mM PMSF) and phosphatase inhibitors (1 mM Na_3_VO_4_ and 1 mM NaF) (Sigma-Aldrich, Saint Louis, MO, USA) for 10 minutes on ice. Lysates were centrifuged at 13,400 rpm for 10 minutes at 4°C, and protein concentrations were determined using the Bradford method ^57^. Proteins were denatured according to Laemmli ^58^, by adding a denaturing β-mercaptoethanol solution and boiling at 100°C for 10 minutes. Proteins were separated by SDS-PAGE and transferred to nitrocellulose membranes (Merck-Millipore, Billerics, MA, USA) using a semi-dry transfer system (Trans Blot Transfer Medium, Bio-Rad). Membranes were blocked with 5% skim milk in TBS-T (20 mM Tris-HCl pH 7.6, 137 mM NaCl, 0.1% Tween-20) 1 hour at 25°C. Immunoblotting was performed using the following primary antibodies: anti-VHR (DUSP3) (1:1000, BD Biosciences), anti-NPM (1:1000, Sigma-Aldrich), anti-p53 (1:1000, Santa Cruz or Cell Signaling) and anti-Actin (1:1000, Santa Cruz). Secondary antibodies conjugated to fluorophores (1:15000, LI-COR Biosciences) diluted in TBS-T were used for subsequent membrane scanning in the Odyssey system (LI-COR Biosciences) using the 683/710 nm or 778/795 nm wavelengths.

### 2.4. Immunofluorescence

Approximately 1x10^6^ cells were collected, centrifuged at 1,000 rpm for 5 minutes to remove the culture medium, washed with PBS and centrifuged again. Cells were fixed for 10 minutes at room temperature (RT) (3% PFA and 2% sucrose in PBS), permeabilized for 5 minutes at 4°C (0.5% Triton X-100, 6.84% sucrose, 3 mM MgCl2 in PBS), blocked for 30 minutes at RT (3% BSA, 10% SFB in PBS) and incubated with the following primary antibodies: anti-DUSP3 (1:500, BD Biosciences), anti-DUSP3 (1:50, Cell Signaling), anti-p53 (1:300, Santa Cruz), or anti-NPM (1:500, Sigma-Aldrich). After primary antibody incubation, cells were incubated with secondary antibodies conjugated to fluorophores Alexa Fluor 488, 555, or 647 (1:300), and resuspended in 4 µL VectaShield (Vector Labs). Confocal microscopy (Zeiss LSM 780) or widefield fluorescence microscopy (Leica DMi8) was used for image acquisition, and images were analyzed using Zen 2.3 (Zeiss) or LAS X (Leica Microsystems AG).

### 2.5. DUSP3 knockdown

DUSP3 was silenced using the GIPZ lentiviral shRNAmir kit (Dharmacon, a Horizon Discovery Group), containing three plasmids targeting different DUSP3 mRNA sequences (S1: AGGTTATAGCCGCTCCCCA; S2: AGGTCCTTCATGCACGTCA; S3: AACGACACACAGGAGTTCA). Lentiviral transduction was performed in RPMI medium without FBS or antibiotics in the presence of 8 μg/mL polybrene. Three MOIs (0.3, 1.0, 3.0) were tested for 2×10D cells, and spinoculation was applied to enhance viral-cell contact. After 72 hours, clonal selection was initiated using puromycin (0.75-2.5 μg/mL). A Mock clone (NS) was created using a non-targeting sequence. Successfully silenced clones were sorted by FACS (BD Aria III) to obtain a homogeneous population expressing GFP and silenced for DUSP3.

### 2.6. Flow cytometry analysis

Approximately 1x10^6^ cells were collected from each condition (Parental, Mock and shRNA DUSP3). Undifferentiated cells were collected by centrifugation, while macrophage-differentiated cells were mechanically detached prior to the centrifugation step. Cells were blocked with 1% BSA for 5 minutes, incubated with the primary antibodies anti-CD14 (1:600) (SC-58951), anti-CD163 (Biolegend 333602), and anti-CD206 (Ab8918) (1:100), following incubation with Alexa Fluor 647-conjugated antibody. Conjugated antibodies PE-anti-CD11b (1:600) (BD555388), PerCP.Cy5.5-anti-HLA-DR (1:500) (BD552764), APC-anti-CD86 (1:500) (BD552764), and APC-anti-CD80 (1:100) (MACS 130-117,719) were also used. Samples were analyzed using the Accuri ™ flow cytometer (BD Biosciences) and BD Sampler software (BD Biosciences).

### 2.7. Data Mining and Bioinformatics

Kaplan-Meyer curves for patient survival analyses were performed correlating the expression of *npm1* and *dusp3* genes in human samples of acute myeloid leukemia (AML). Genetic correlations were analyzed in whole blood from CML and AML patients using the GEPIA2 website (gepia2.cancer-pku.cn) with data from the TCGA Tumor (LAML = AML -Acute Myeloid Leukemia) and GTEx (Whole Blood and Chronic Myelogenous Leukemia - CML) databases (accessed 09-06-2024).

### 2.8. Statistical Analysis

Statistical analyses were performed using ANOVA followed by Tukey’s post-test. Results were obtained from independent triplicates and expressed as mean ± SD. Graphical representation and statistical analysis were performed using GraphPad Prism 8.0. Significance levels: *p<0.05; **p<0.01; ***p<0.001; \*\*\**p<0.0001.* Asterisks (*) represent significance between parental and transduced cells, and hashes (#) represent significance between Mock and transduced cells.

## 3. RESULTS

### DUSP3 is differentially expressed in acute myeloid leukemia cells

DUSP3 has been described as a regulatory protein for several cellular processes, such as proliferation, differentiation, ^18^ and DNA repair ^19^. In this context, changes in its expression and localization in cells at different stages of differentiation are potentially expected. To explore this further, we performed western blotting (Fig. 1A and 1C) and immunofluorescence (Fig. S1) to assess DUSP3 levels in myeloid parental HL-60 and THP-1 cells. In both experiments, we observed that DUSP3 expression is higher in THP-1 than HL-60 cells. We also examined the basal expression of DUSP3, NPM and p53 in both cells, in the absence and presence of differentiation stimulus to macrophages using PMA, by western blotting (Fig. 1B) and immunofluorescence (Fig. S2). The MRC-5 cell line (human normal lung fibroblasts) was used as a positive control for protein expression (Fig. 1B). NPM was present in the nucleus of both cell lines (Fig. S2) and differentiation into macrophages did not change the NPM expression in THP-1 cells, though a slight increase in DUSP3 expression was observed (Fig. 1D). In differentiated HL-60 cells a marked decrease in the levels of DUSP3 and NPM was detected (Fig. 1D). Notably, in both cell lines there was no detectable expression of p53 protein, despite the differentiation stimulus (Fig. 1, S1 and S2B). Bioinformatics data support these findings, showing the different expression of DUSP3 between these two cell lines (Fig. S3A).

**Figure 1.**
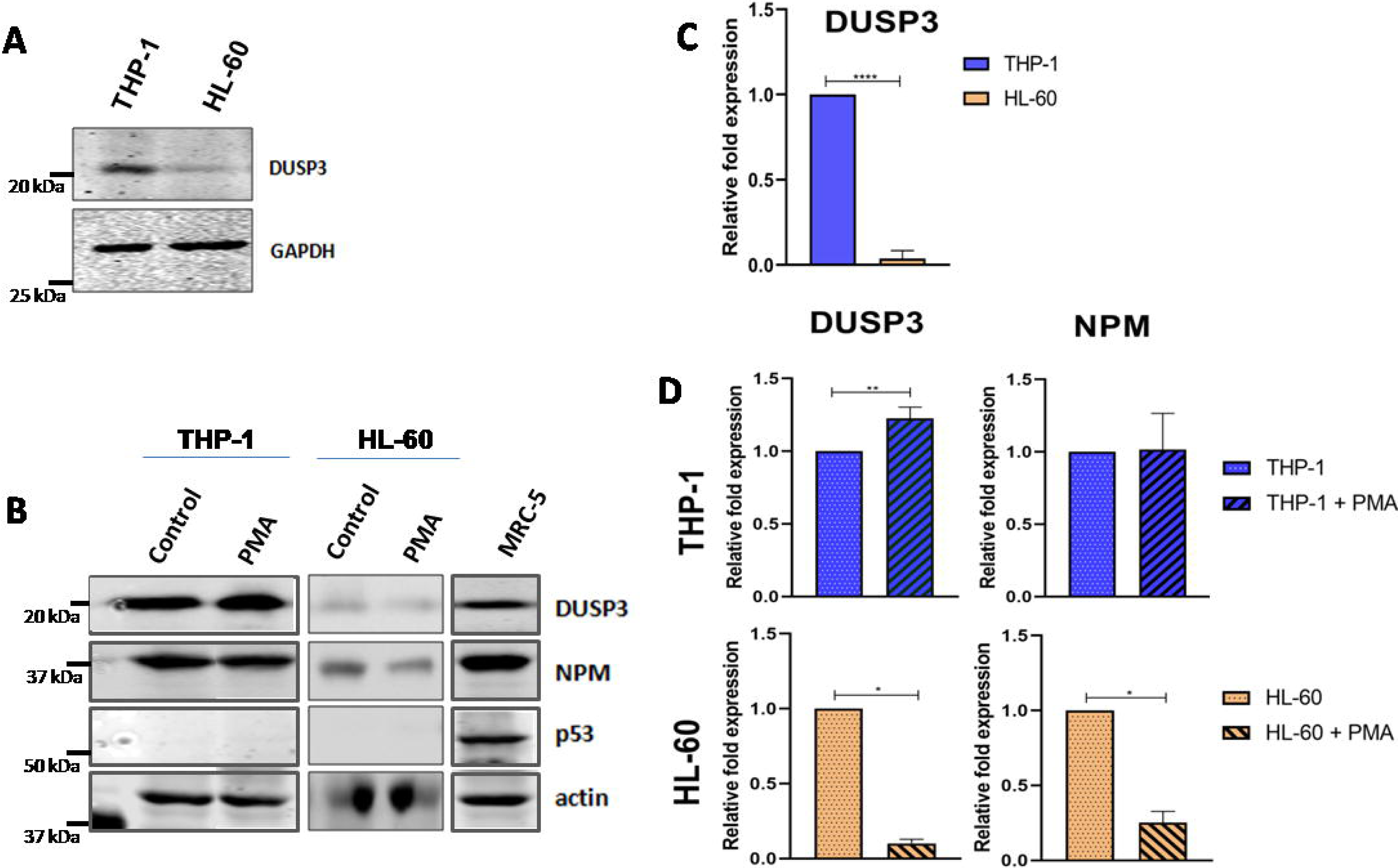
DUSP3 is differentially expressed during acute myeloid leukemia cells maturation. (A) Basal levels of DUSP3 expression differ between untreated, undifferentiated THP-1 and HL-60 cells. (B) Upon PMA-induced differentiation into macrophages, differences in DUSP3 and NPM expression were observed, while p53 expression remained unchanged. (C, D) Quantification of the Western blots from panels (A) and (B), respectively, was performed by densitometry using Image Studio Lite Ver 5.2. DUSP3 and NPM band intensities were normalized to GAPDH or Actin. Data are represented as mean ± SD.

### 3.1. The DUSP3 knockdown contributes to the modulation of NPM expression

After observing that differentiation changed DUSP3 and NPM protein levels, we proceeded with the DUSP3 knockdown by stably transducing THP-1 and HL-60 cells with three different shRNA sequences targeting the *dusp3* gene (Fig. 2). The cell population silenced for DUSP3 was selected by the GFP-tag present in the lentiviral vector construct through FACS, and GFP-related fluorescence was also monitored by light microscopy (Figs. S4 and S5). The nomenclature of clones was given as follows: the lentiviral vector with a shRNA that does not match any other present in the cell was called Mock; vectors containing shRNA for DUSP3 received a prefix "S" (for silenced), and the three different sequences were named from 1 to 3. For THP-1 cells, different MOIs (multiplicity of infection) were tested, which explains the number after the identification of the sequence of shRNA used. In the case of HL-60 cells, an MOI of 1 was optimal for all transductions, so no second number is included in this clone’s nomenclature. After knockdown, the expression levels of DUSP3 and NPM proteins were monitored in three isolated clones for each cell line (Fig. 2A). Parental cells were used as a non-transduced control, while Mock was used as a control of the transduction process to account for any cellular perturbations. In both cell lines, at least one of the three different shRNA sequences for DUSP3 achieved over 50% knockdown of DUSP3 protein levels. As shown in Fig 1B, where HL-60 differentiation caused a decrease in the levels of DUSP3 and NPM levels, DUSP3 knockdown also led to a reduction in NPM expression (Fig. 2B). A direct and expected effect of DUSP3 knockdown was also observed for the ERK1/2 phosphorylation in THP-1 and HL-60 cells and clones (Fig. S6).

**Figure 2.**
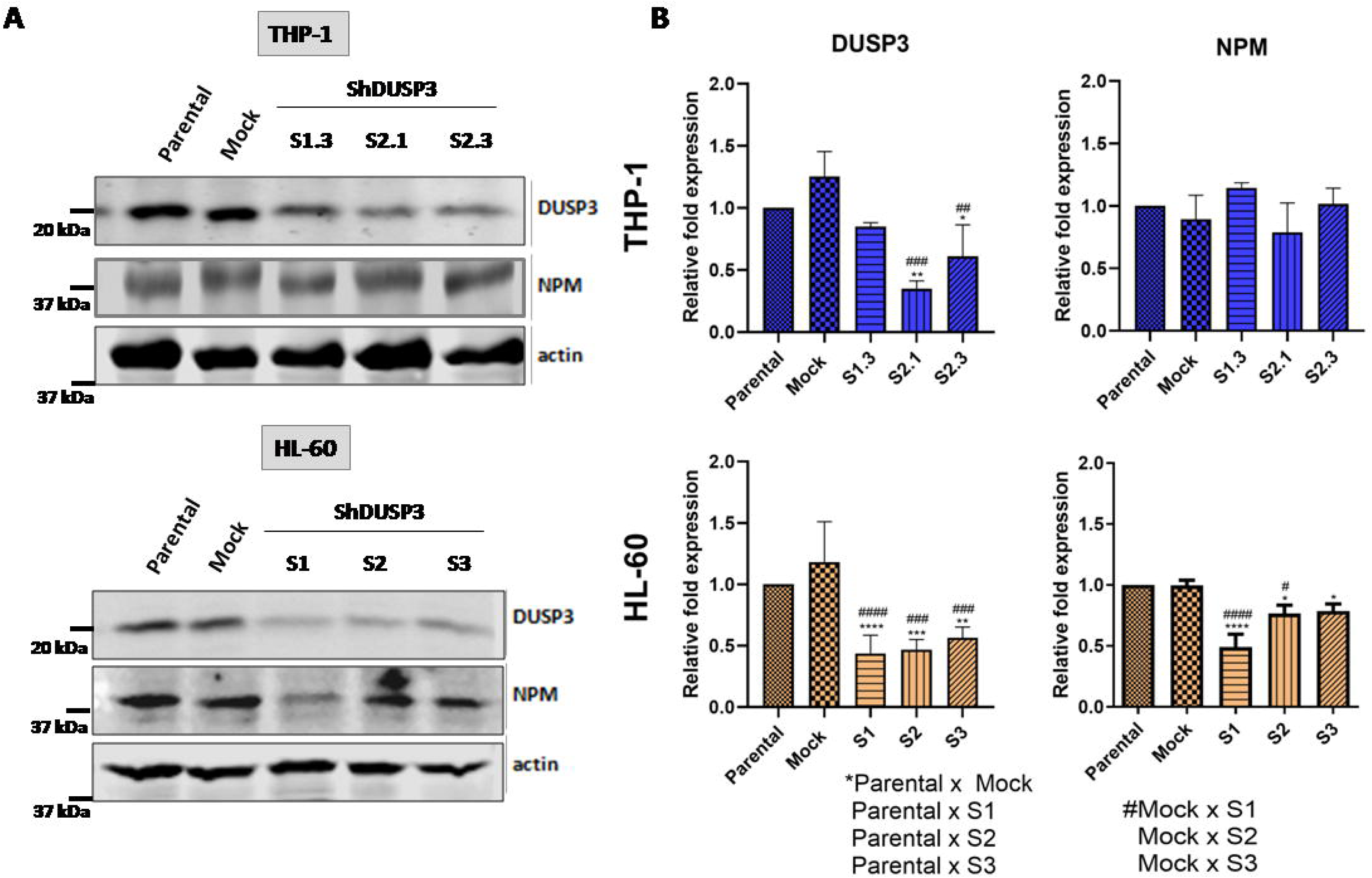
DUSP3 knockdown modulates NPM expression. (A) The effectiveness of DUSP3 knockdown was verified by Western blot in both cell lines, as well as its impact on NPM expression. (B) Densitometric analysis was performed by Image Studio Lite Ver 5.2, with DUSP3 and NPM bands normalized to Actin. Data are represented as mean ± SD.

### 3.2. DUSP3 knockdown contributes to the differentiation of acute myelocytic leukemia cells

To verify a possible involvement of DUSP3 knockdown in the differentiation of myeloid cells into macrophages, we performed flow cytometry analysis to investigate markers commonly altered during myeloid cells differentiation. We initially assessed the expression of CD11b and CD14 proteins in THP-1 and HL-60 cells (Fig. 3A). For this purpose, we evaluated the most efficiently silenced shDUSP3 clones (THP-1 S2.1 and HL-60 S1) and compared them with the parental cells and Mock clones, before and after treatment with PMA to induce differentiation into macrophages. In undifferentiated THP-1 cells no significant differences in CD14 and CD11b expression were observed between the DUSP3-silenced clone S2.1 and controls. However, in differentiated THP-1 cells, DUSP3 knockdown (THP-1 S2.1) led to a ∼40% reduction in CD14 levels (Fig. 3A upper), with no significant changes in CD11b expression. In HL-60 cells, regardless of the differentiation stimulus, DUSP3 knockdown resulted in a ∼50% decrease in CD14 expression (HL-60 S1 clone). In the case of undifferentiated HL-60 cells, an increase of 60% in CD11b expression was observed in clone S1 compared to controls (Fig. 3A lower). Notably, CD11b levels increased in both cell lines following PMA stimulation, although they were slightly lower in the DUSP3-silenced cells compared to Mock clones, as the transduction process itself caused an increase in CD11b levels. Once again, the bioinformatics data revealed altered expression of genes involved in the differentiation pathway between the HL-60 and THP-1 lines, corroborating the experimental data (Fig. S3B).

**Figure 3.**
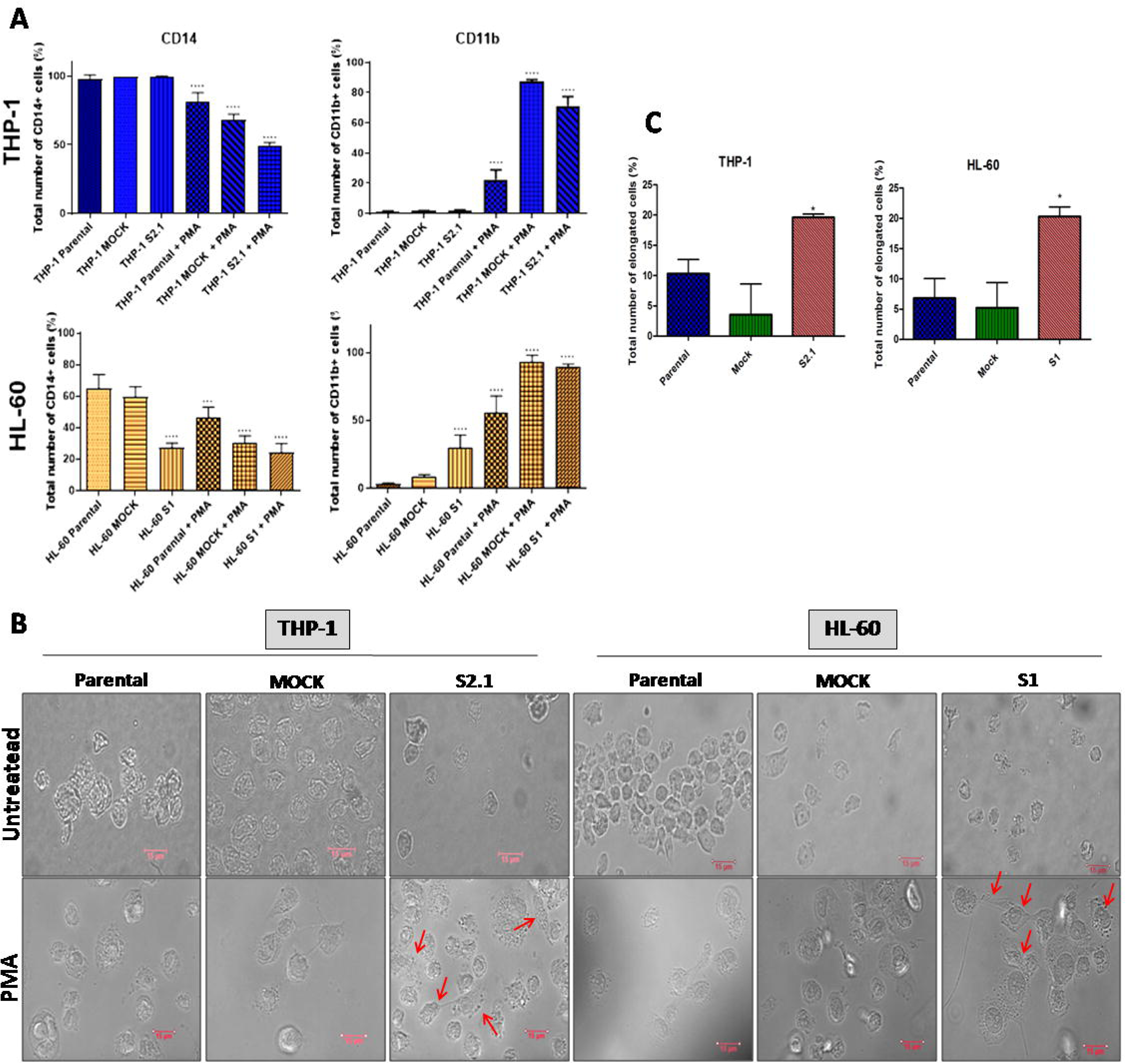
DUSP3 knockdown promotes differentiation in acute leukemic cells. (A) DUSP3 knockdown modulated the expression of the differentiation markers CD14 and CD11b. In silenced cells without PMA treatment, this modulation was evident only in HL-60 cells, where CD14 expression decreased and CD11b expression increased. (B) THP-1 and HL-60 clones were treated with PMA for differentiation into macrophages, compared to their respective differentiated controls (Parental and Mock), as visualized in bright-field microscopy. Red arrows indicate cells with elongated morphology following PMA treatment. (C) Quantification of elongated cells post-differentiation. Data are presented as mean ± SD.

### 3.3. DUSP3 knockdown contributes to macrophages polarization

To investigate a potential cellular polarization phenotype resulting from the THP-1 and HL-60 cells differentiation following PMA stimulus, as well as the influence of DUSP3, we examined the morphology of these cells using brightfield microscopy (Fig. 3B). Corroborating the expression of differentiation markers, DUSP3 knockdown alone induced morphological changes, since it was possible to observe the emergence of a mixed population containing elongated cells, suggesting an alternative cellular phenotype ^22^ (Fig. 3B). The quantification of these elongated cells revealed a significant enrichment of this population in both DUSP3-silenced clones (Fig. 3C). Next, we performed flow cytometry analyses to assess surface protein markers expressed in classically activated macrophages (M1 phenotype) and alternatively activated macrophages (M2 phenotype). Initially, we examined three markers commonly associated with M1-like macrophages: CD80, CD86, and MHCII ^23^. In both cell lines, regardless of the PMA differentiation stimulus, no significant differences were found in CD80 and CD86 expression between control cells and DUSP3 knockdown clones. For MHCII, no changes were detected in undifferentiated cells when comparing controls to DUSP3-silenced clones, but in Mock clones, PMA stimulation led to a substantial increase in MHCII expression, which was reduced by DUSP3 silencing (Fig. 4). As these results were suggestive of a macrophage polarization into the M2-like phenotype, we analyzed two additional markers, CD163 and CD206, which are commonly used to identify the alternative activation of macrophages^24^ (Fig 5). Contrary to expectations, no changes in CD163 and CD206 expression were detected in undifferentiated cells from either cell line, with both markers displaying low levels. After PMA-induced differentiation, the CD163 expression was markedly elevated, while CD206 remained relatively low. In contrast, DUSP3 knockdown resulted in the opposite trend, with a significant reduction in CD163 and a pronounced increase in CD206 compared to the two control conditions. Collectively, these results indicated that DUSP3 knockdown, combined with PMA differentiation stimuli, drives THP-1 and HL-60 cells toward an alternative macrophage activation phenotype (M2-like).

**Figure 4.**
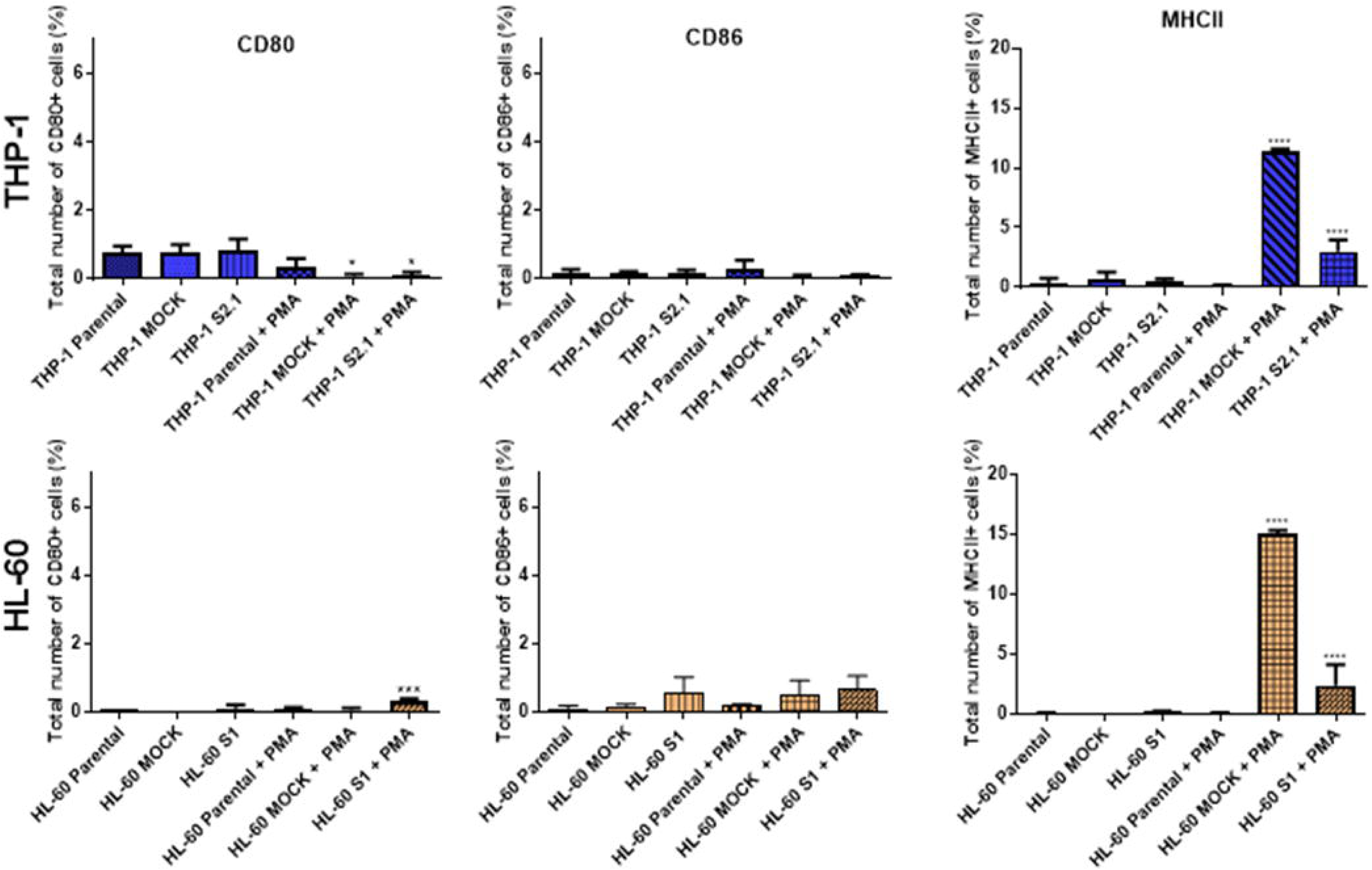
DUSP3 knockdown is associated with reduced M1-like macrophage markers in acute myeloid leukemia cells. DUSP3 knockdown did not result in significant changes in CD80 and CD86 expression, regardless of PMA treatment. However, for MHCII, only cells transduced with the non-silencing control vector (MOCK) exhibited an increase in expression after PMA treatment compared to Parental cells, while DUSP3 silencing led to a reduction in this marker. Data are presented as mean ± SD.

**Figure 5.**
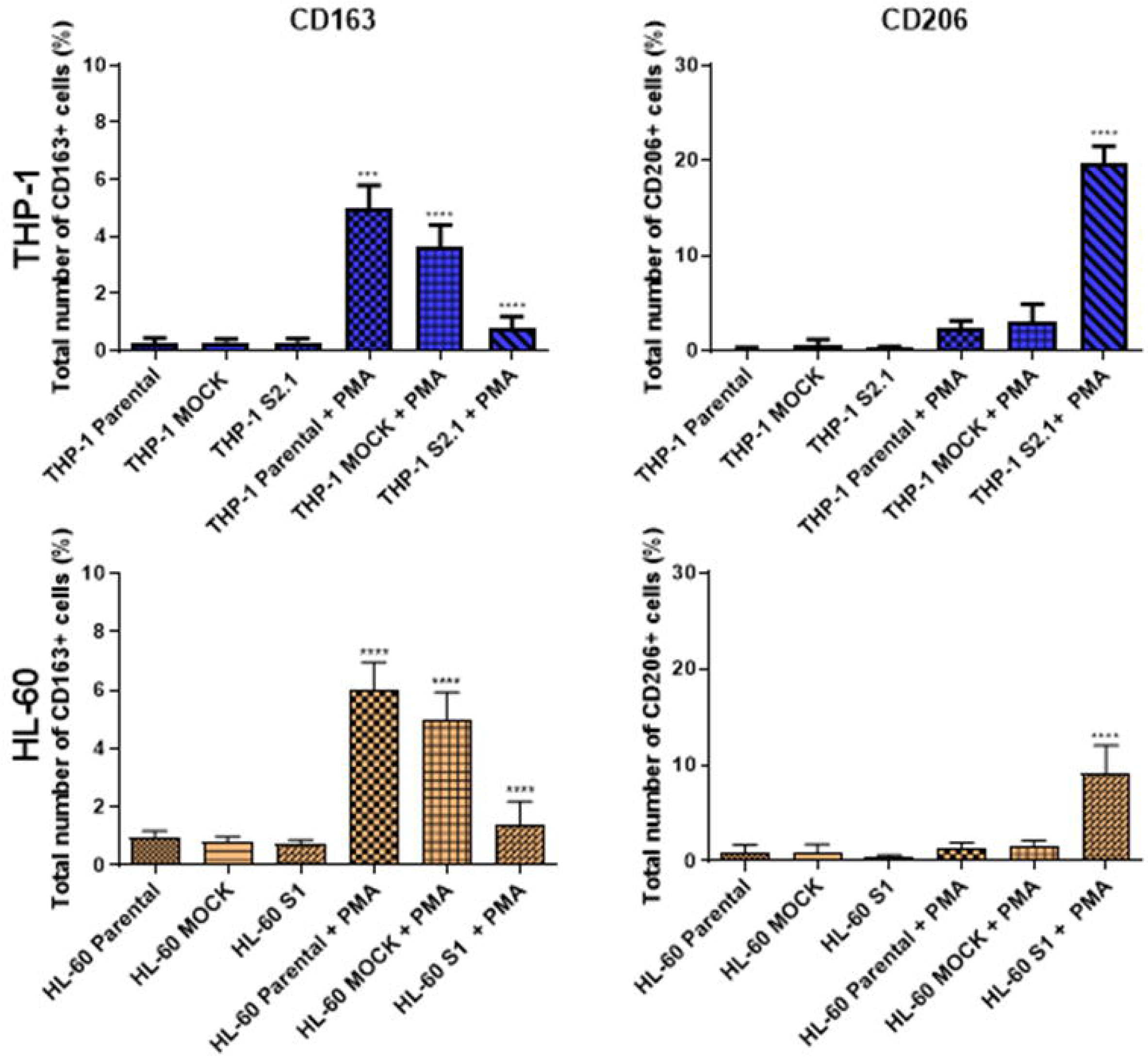
DUSP3 knockdown increases M2-like macrophage markers in acute myeloid leukemia cells. The DUSP3 knockdown did not significantly alter CD163 and CD206 expression in the absence of PMA treatment. However, following PMA-induced differentiation, CD163 expression increased significantly in Parental and Mock clones but decreased in DUSP3-silenced clones for both cell lines. CD206 expression increased significantly after differentiation in silenced clones. Data are presented as mean ± SD.

### 3.4. The *dusp3* gene expression is related to the survival of patients with acute myeloid leukemia and may be associated with the affected pathways

Kaplan-Meier curves were used to assess AML patient survival in relation to the expression of *dusp3* and *npm1* genes, normalized by the *actb* gene expression (Fig. 6A). For *dusp3*, we found that within approximately 2.5 years (up to 30 months) its expression does not seem to affect survival; however, after this period, there is a shift in this trend, with patients exhibiting high *dusp3* expression showing reduced survival. In the case of *npm1*, the lower expression is directly correlated to a markedly low patient survival up to approximately 4 years, after which any correlation between *npm1* expression and survival is lost. Nonetheless, in both undifferentiated myeloid cells subjected to DUSP3 knockdown, no significant differences in cell proliferation were observed compared to Mock cells (Fig. S7), even in the HL-60 clones where NPM levels are indirectly reduced by the DUSP3 silencing (Fig. 2A). Enrichment analysis of 500 genes essential for overall survival in AML revealed that the most affected pathways include DNA repair, differentiation, cell cycle, NF-kB, p53, as well as pathways regulated by MAPKs and tyrosine phosphatases (Fig. 6B). Differentially expressed genes in AML were also evaluated, correlating with these same observed pathways that affect overall survival (Fig. 6C).

**Figure 6.**
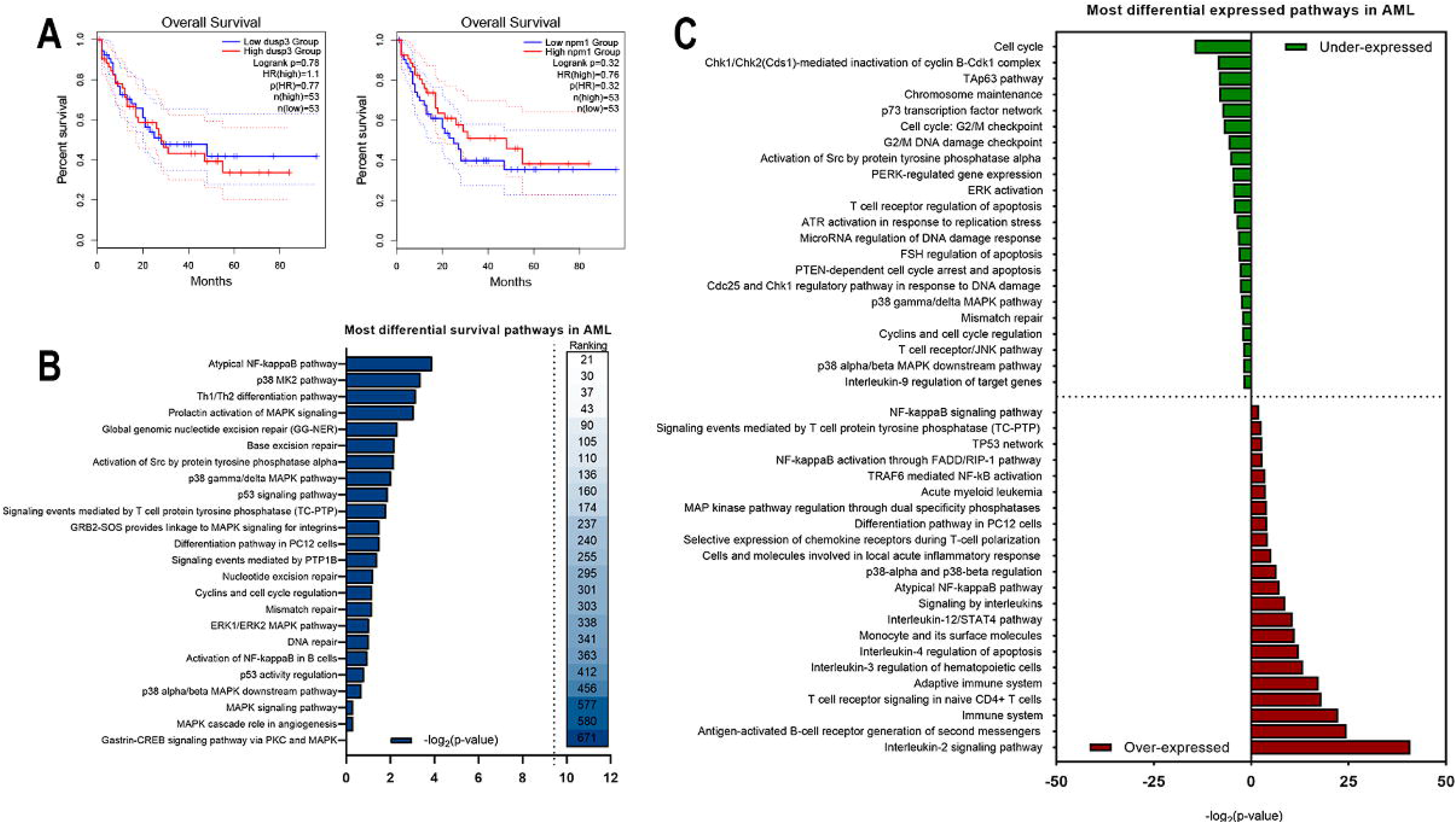
DUSP3 gene expression correlates with acute myeloid leukemia patient survival, with MAPKs also influencing overall survival. Over an average of 5 years, *dusp3* gene expression did not significantly impact the survival of patients with acute myeloid leukemia (AML). However, high *npm1* expression over the same period appeared to favor patient survival. In shorter periods, such as approximately one year, the effect of *dusp3* expression was less pronounced, while higher *npm1* expression was linked to increased survival.

### 3.5. The expression of *dusp3* and *npm1* genes shows correlation with both blood cell maturation and expression of genes encoding markers of cell differentiation

To explore a possible correlation between DUSP3 and NPM proteins, gene expression analyses were conducted in whole blood, chronic myelogenous leukemia (CML) and acute myeloid leukemia (AML) using the GEPIA2 database (Fig. 7). A positive correlation between the *dusp3* and *npm1* genes was observed in CML, which is rich in mature cells, as indicated by Pearson’s correlation coefficient (r). This correlation weakens in samples with lower degrees of cell maturation, such as whole blood and AML (Fig. 7A). Furthermore, correlations between the *dusp3* and *npm1* genes and the differentiation markers CD14 (Fig. 7B-C) and CD11b (Fig. 7D-E) were observed. For example, *dusp3* expression influences the *cd14* expression according to the degree of cell maturation: the more differentiated the cell, the stronger the positive correlation, which tends to decrease with cell immaturity, as observed in AML (Fig. 7B). The correlation between *npm1* and *cd14* genes is lower in CML and AML and almost non-existent in whole blood, where the Pearson correlation coefficient approaches zero (Fig. 7C). The correlations for *cd11b* followed a similar trend to *cd14*: a positive correlation with *dusp3* expression in more mature cells, which weakens in AML and even tends toward a negative correlation, where high expression of one gene coincides with reduced expression of the other (Fig. 7D). Similarly, the correlation between *npm1* and *cd11b* is slightly stronger in CML samples and tends toward a negative correlation in AML, indicating a possible relationship between *dusp3*, *npm1*, and the degree of cell differentiation (Fig. 7E).

**Figure 7.**
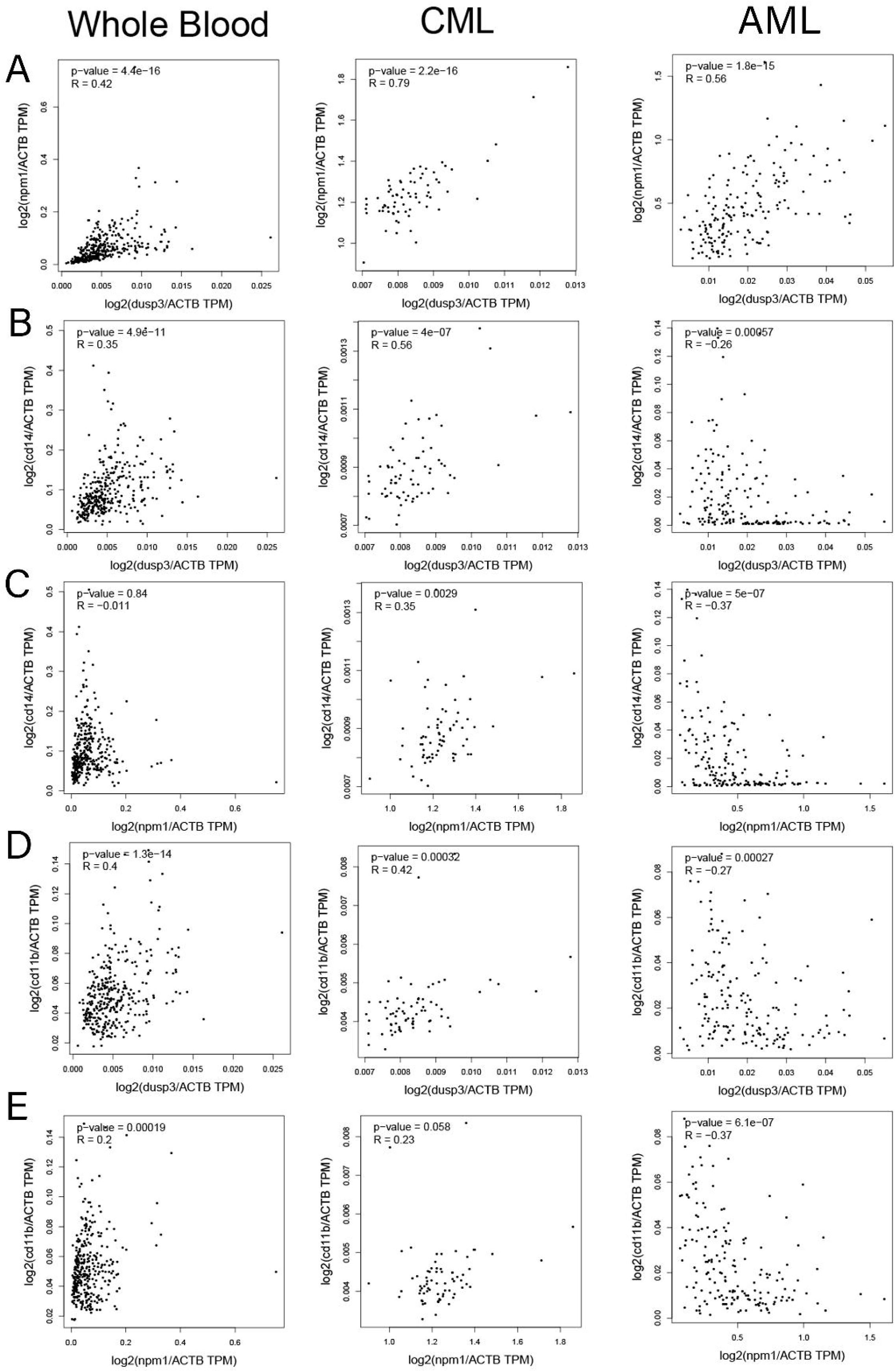
Correlation analysis of *dusp3, npm1, cd14, and cd11b* gene expression. (A) In whole blood samples, which consist of various cells at different stages of maturation, there is a progressively higher positive correlation between *dusp3* and *npm1* expression as cells mature, such as in CML. (B) The correlation between *dusp3* and *cd14* expression is positive in mature blood cells but decreases with cell immaturity (whole blood and AML, respectively). (C) *npm1* shows a weaker correlation with *cd14* in mature cells, which becomes even weaker and negative in whole blood and AML. (D) *dusp3* correlates positively with *cd11b* expression in mature blood cells, but this correlation diminishes with increasing cell immaturity in AML. (E) The correlation between *npm1* and *cd11b* expression is positive in mature cells but turns negative in AML.

### 3.6. The *npm1* gene expression correlates with the p21^Cip1^ and NF-_D_B pathway genes in blood mature cells

To explore potential interrelations between DUSP3/NPM with mechanisms of cell differentiation and proliferation found in this work, we analyzed Pearson’s correlation for the gene pairs *dusp3* x *cdkn1a* and *npm1* x *cdkn1a*, as the *cdkn1a* gene encodes the well-known cell cycle inhibitor p21^Cip1^ (Fig. 8A-B). A positive correlation was observed between the *npm1* x *cdkn1a* genes in whole blood and CML, which became negative in AML, while for *dusp3* this correlation remained positive across different leukemia types, although with a tendency to decrease in AML. We also investigated a possible correlation between *dusp3* and *npm1* genes with the transcription factor NF-ĸB (Fig. 8 C-F), a heterodimer composed of p65 (*rela* gene) and p50 (*nfkb1* gene) protein subunits, which binds to a response element within the *cdkn1a* gene promoter to initiate its transcription. Both *dusp3* and *npm1* genes showed a positive correlation with the *rela* and *nfkb1* genes in whole blood cells and AML. In CML, the correlation weakened for *rela* but strengthened for *nfkb1*, consistent with the higher degree of cellular maturity. These correlations suggest that NF-ĸB pathway may serve as a link between DUSP3-NPM and p21^Cip1^ for the differentiation and maturation of myeloid cells despite their proliferation. Supporting these observations, we found a strong Pearson’s correlation between dusp3 or *npm1* and *tnf-*_LJ_ genes in more mature blood cells, but not in immature AML cells (Fig. S8). These findings align with literature reports showing that TNF-α treatment promotes NPM oligomerization, very likely due to the NPM phosphorylation at Ser125 by IKKα, followed by the translocation of p65 NF-ĸB subunit into the nucleus ^25^. It was demonstrated by co-immunoprecipitation that NPM interacts with the p65 subunit, with this interaction increasing after NF-ĸB activation by TNF-L ^26^. Thus, NPM and NF-ĸB act coordinately to regulate gene transcription broadly, including inflammatory cytokine genes as TNF-L ^27^.

**Figure 8.**
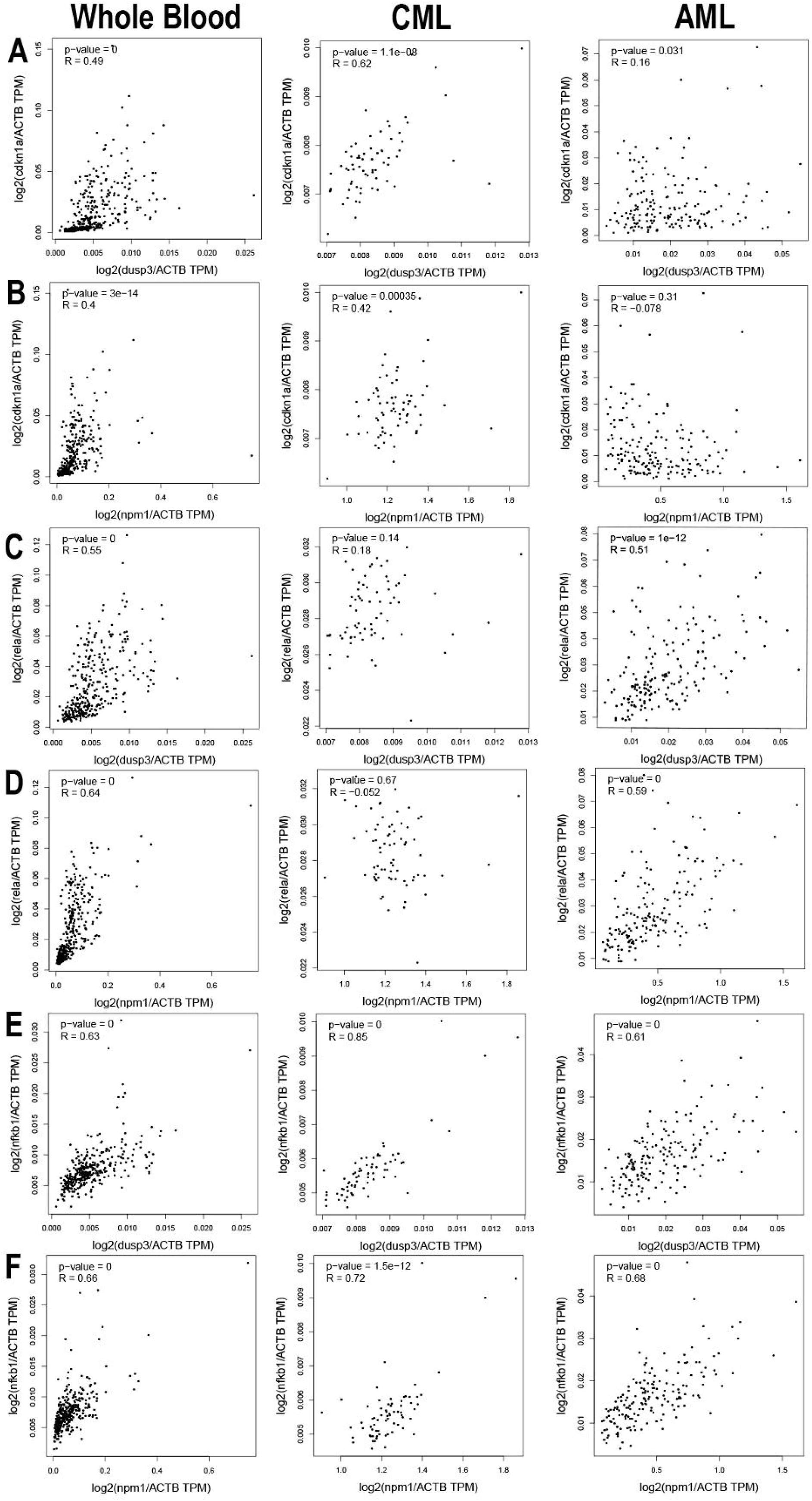
Correlation analysis of *cdkn1a*, *rela* and *nfkb1* gene expression. (A) A positive correlation exists between *dusp3* and *cdkn1a* expression, which decreases with increasing cell immaturity (whole blood and AML, respectively). (B) *npm1* shows a positive correlation in whole blood and CML but becomes negative in AML cells. (C) *dusp3* correlates positively with *rela* expression in whole blood and AML, decreasing in CML. (D) *npm1* shows a negative correlation in CML, while in whole blood and AML, this correlation becomes positive. (E) The correlation between *dusp3* and *nfkb1* is positive, strongest in CML, and decreases in AML and whole blood. (F) *npm1* shows a positive correlation with *nfkb1* in whole blood, CML, and AML, though less than of *dusp3* in CML (E).

## 4. DISCUSSSION

The cell lines THP-1 ^28^, derived from acute myeloid leukemia (AML), and HL-60 ^29^, derived from acute promyelocytic leukemia (APL), express the DUSP3 enzyme at different levels and cellular localizations (Fig. 1A, S1 and S2). THP-1 cells show the highest levels of DUSP3 and NPM, both localized in the nucleus, with NPM being predominantly nucleolar (Fig. S2). Since these two leukemic cells have different degrees of maturation and DUSP3 expression, we sought to induce their differentiation into macrophages through PMA stimulation, as the *dusp3* gene is highly expressed in macrophages ^12^ (Fig 1B). Unlike THP-1 cells, the differentiation of HL-60 into macrophages resulted in a decrease in the levels of DUSP3 and NPM (Fig. 1B), which was contrary to expectations, as primary macrophages are reported to have high levels of this phosphatase compared to other immune cells, such as neutrophils, T cells, and B cells ^30^. Genetic correlations extracted from patient data indicate that AML exhibits a weaker correlation between *dusp3* and *npm1* expression. However, when extrapolated to CML samples, which contain a higher proportion of mature cells, this correlation tends to become positive and significant (Fig. 7A). A modest increase in DUSP3 expression, but not in NPM, was observed 96 hours after PMA stimulation in THP-1 cells and was associated with a significant decrease in ERK1/2 phosphorylation (Fig. S9). MAPKs are well-known substrates of DUSP3 ^17^, and it has been shown that PMA, a classical MAPK activator, induces the expression of the Pyst2-L phosphatase in leukemic cells, suggesting a role for MAPK-Pyst2-L signaling pathway in these malignant processes ^14^. DUSP5 is also transiently induced in myeloid cells by differentiation stimuli, and its knockdown affects the differentiation of myeloid progenitors into macrophages after stimulation with macrophage colony-stimulation factor (M-CSF) stimulation, through sustained activation in ERK1/2^15^.

We generated DUSP3-silenced THP-1 and HL-60 cells, with at least one sequence of shDUSP3 achieving approximately 50% knockdown of DUSP3 protein levels, and we evaluated changes in NPM levels. In HL-60 cells, DUSP3 knockdown alone caused a significant decrease in NPM levels (Fig. 2), while no changes were observed in THP-1 cells. The maintenance of NPM expression in THP-1 cells is probably related to the partial silencing of DUSP3 (∼50%) since this cells already expresses high basal levels of this protein. In contrast to what have been reported for DUSP5, DUSP3 knockdown did not significantly affect ERK1/2 phosphorylation in both cell lines (Fig. S5) ^15^, suggesting a DUSP3-independent mechanism of ERK1/2 regulation. According to these data, we propose that in HL-60 cells the DUSP3 knockdown is somehow correlated with the decrease in NPM protein levels (Fig. 2), and this effect is also observed during HL-60 differentiation into macrophages following PMA stimulation (Fig. 1B). Alterations in NPM levels are closely linked to leukemogenesis, since its overexpression has been detected in leukemia blasts and cell lines ^31,32^. High levels of NPM protein have been implicated in diminishing the susceptibility of human leukemic HL-60 cells to retinoic acid (RA)-induced apoptosis ^33^, and promoting resistance to differentiation of K562 cells induced by either RA or PMA ^34^. On the other hand, NPM knockdown in HL-60 ^35^ and K562 ^36^ cells resulted in decreased cell proliferation, blockade of the cell cycle progression and increased expression of pro-apoptotic genes and proteins. Interestingly, NPM knockdown in HL-60 cells also induces differentiation ^35^. Although we did not observe significant differences in HL-60 proliferation after DUSP3 knockdown (Fig. S6), this phenotype promoted a decrease in NPM levels and notable differences in the expression levels of differentiation markers CD14 and CD11b, as determined by flow cytometry in both myeloid cell lines (Fig. 3A).

CD14 is a polysaccharide-binding protein highly expressed in blood monocytes and tissue macrophages, and despite poorly expressed in neutrophils, induced cellular maturation can increase or decrease its levels ^37^. CD11b is mostly expressed by granulocytes, monocytes, macrophages, and NK cells, among other immune cells ^38^. Immunophenotyping assays for CD14 and CD11b revealed that PMA-induced differentiation of both THP-1 and HL-60 cells into macrophages is characterized by a reduction in CD14 expression and an increase in CD11b expression (Fig 3A). In HL-60, DUSP3 silencing alone seems to behave as an inducer of cell differentiation, as evidenced by the decrease in CD14 expression and the increase in CD11b expression, even in the absence of PMA stimulation, which was not observed for THP-1 cells. In agreement with that, gene expression of *dusp3* is positively correlated with *cd14* and *cd11b* in more mature cells, whereas *npm1* shows a weaker correlation with these markers (Fig. 7B-E).

Macrophage activation is an emerging area of immunology, tissue homeostasis, disease pathogenesis, and control of inflammatory processes ^39^. In this work, we used the terms M1 and M2 macrophages as synonyms for classical activation and alternative activation, respectively. According to our cellular morphology data, DUSP3 knockdown cells, following PMA treatment, acquired a mixed morphology, with the appearance of elongated cells (Fig. 3B and 3C). This morphological change suggested that these cells might express lower levels of inflammatory markers typically associated with M1-type macrophages. CD80 and CD86 markers are co-stimulatory molecules involved in the regulation of inflammation and tissue damage in M1-polarized macrophages. At baseline, macrophages express low levels of CD80 and constitutively express CD86, but both molecules can have their expression induced after stimulation with LPS or IFN-γ ^40,41^. However, our immunophenotyping assays revealed that the expression of these markers was not significantly altered in either cell lines, regardless of the absence or presence of differentiation stimulus, which highlights the fact that they are not related to M1 phenotypes (Fig. 4). MHCII receptor, another important marker of M1 macrophages, can also be present at low levels in M2 macrophages^42^. In this study, MHCII expression was detected in both undifferentiated and PMA-treated cells. Notably, transduced cells (Mock and shDUSP3 clones) treated with PMA exhibited a significant, but unexpected, increase in MHCII expression in Mock cells and a decrease in shDUSP3 cells. Therefore, these data suggest that DUSP3 silencing is not associated with the M1 phenotype (Fig. 4).

M2 macrophages are defined by their functional properties, including the high expression of the mannose receptor (CD206), IL-10, and angiogenic factors, along with low expression of IL-12, and its involvement in immune regulation and tissue remodeling ^43,44^. The hemoglobin scavenger CD163 is a specific marker for alternatively activated (M2-type) macrophages and is responsible for the uptake of hemoglobin released into the plasma. Its expression is regulated during inflammation, making CD163 a hallmark of tissue inflammation ^45^. In our study, no changes in the levels of these two markers were observed in THP-1 and HL-60 cells without PMA treatment. However, following PMA-induced differentiation into macrophages, DUSP3 knockdown resulted in a decrease in CD163 expression in both cell lines, which may be related to the absence of inflammatory stimuli, such as cytokines (Fig. 5). For the CD206 marker, a receptor widely recognized for its presence in M2 macrophages and immature dendritic cells ^46^, we detected a significant increase in its expression in both myeloid cells treated with PMA, particularly in the DUSP3-silenced cells. This finding suggests a strong association between DUSP3 knockdown and M2 macrophage polarization (Fig. 5).

A previous study demonstrated that DUSP3-deficient mice exhibit tolerance to LPS-induced endotoxic shock and polymicrobial septic shock. This was characterized by a marked increase in anti-inflammatory M2-type macrophages, decreased TNF-L production, and reduced ERK1/2 activity, with no changes in T/B lymphocytes, neutrophils, monocytes, or platelet counts ^47^. Subsequently, it was shown that DUSP3’s involvement in this immune response is influenced by sex hormones produced by the host animals. The resistance to sepsis of female mice (except DUSP3^-/-^) was associated with reduced activation of ERK1/2, PI3K, and AKT pathways, likely due to a deficiency in phosphatase activity. These findings suggest that, in the absence of DUSP3, female sex hormones are transcriptionally implicated in the acquired resistance of DUSP3 ^-/-^ mice to LPS-induced lethality ^30^. In contrast, we observed here in two human acute myeloid leukemia cell models with different maturation stages, that DUSP3 knockdown is involved with the differentiation and polarization of these cells into M2-type macrophages through mechanisms that do not necessarily involve ERK1/2, p53, or *in vivo* hormonal action systems. Other studies have also shown that the inhibition of DUSP3 activity in THP-1 cells induced by LPS induces a shift from M1 to M2 polarization ^48^. Additionally, it has been demonstrated that the regulatory pathway involved in the M2 phenotype, particularly in association with other tumor lineages, is mediated through the DUSP3/JAK2/STAT3 axis. This refined control of pro-tumor activity may enhance the potential for tumor infiltration and metastasis ^49^.

DUSP3 knockdown in HL-60 cells resulted in decreased NPM expression, which may be associated with decreased TNF-L expression. In contrast, silencing DUSP3 in THP-1 cells did not affect NPM expression. It has been shown that PMA effects on HL-60 cells are TNF-L independent, since PMA does not induce TNF-L synthesis in these cells ^48^. The modest effect of PMA stimulation observed in THP-1 cells may be partially attributable to TNF-L induction, as both TNF-L and PMA are known inducers of the NF-ĸB transcription factor. The involvement of the tumor suppressor gene *tp53* and the p53 protein has been well established in various human cancers, including leukemia and lymphomas. For example, the THP-1 cell line harbors a frameshift mutation caused by a 26-base pair deletion starting at codon 174, resulting in non-recognition by wild-type p53 cDNA ^51^. On the other hand, the HL-60 cell line is p53-null as it does not express detectable levels of p53 protein or mRNA due to large deletions in the *tp53* gene ^52^. When tested for p53 protein expression, neither cell line exhibited detectable or significant levels of p53 (Figs. 1B and S1), which led us to investigate other potential pathways involved in promoting differentiation.

NPM and NF-ĸB act as components of basal the transcription complex, which may influence *cdkn1* gene transcription and potentially negatively regulate inflammatory cytokines ^27^. In that sense, NPM plays dual roles: acting as an NF-κB co-activator to induce *sod2* gene expression ^53^ and serving as a co-repressor during retinoic acid (RA)-induced differentiation by recruiting histone deacetylases (HDACs) in gene expression ^54,55^. Additionally, NPM has histone chaperone activity, contributing to chromatin organization and transcriptional control ^56^. Thus, we can assume that the reduced levels of NPM in HL-60 cells correlates with a lower repression of gene transcription, presumably due to a decreased recruitment of HDACs, resulting in increased transcription of differentiation-related genes. Our experimental data may also be associated with NPM’s reduced role as histone chaperone, affecting chromatin remodeling and global gene transcription. For example, this would explain the changes of CD14 and CD11b markers found in HL-60 cells under DUSP3 knockdown, even in the absence of differentiation stimulus. In contrast, the maintenance of NPM levels in THP-1 cells with DUSP3 knockdown could be linked to stronger repression of inflammatory cytokine transcription, co-activation of NF-ĸB pathway, and increased TNF-L production, which altogether can elevate HDACs activity and further repressing differentiation genes expression. In HL-60 cells under DUSP3 knockdown, the decrease of NPM functions as a co-activator of the NF-ĸB pathway is supported by a strong positive correlation between *npm1 and rela* genes in more mature cells, as well as the *nfkb1* expression being more correlated with *dusp3*, while a decrease in this pathway appears to impact *cdkn1a* gene expression. In THP-1 cells with DUSP3 knockdown, a positive correlation persists between *npm1* and *rela*, but the correlations between *nfkb1* and *dusp3*, as well as *cdkn1a* and *dusp3*, are less pronounced, likely due to TNF-L production maintaining NF-ĸB pathway activation.

## 5. CONCLUSIONS

In summary, this work provides evidence of the DUSP3’s involvement in the differentiation and polarization of myeloid leukemia cells into macrophages, in a maturation stage-dependent manner, likely influenced by changes in NPM functions that affect pro-inflammatory TNF-L signaling. We observed, through bioinformatics analyses, that higher DUSP3 expression correlates with reduced survival rates after 4 years, whereas increased NPM levels are associated with improved patient survival up to approximately 5 years. Thus, by improving the knowledge of these proteins and the signaling pathways they regulate, we propose the DUSP3-NPM axis as potential therapeutic target for leukemia treatment through the interference with cell differentiation processes, potentially reducing resistance to conventional therapies. In addition, the polarization of macrophages toward the M2 phenotype may attenuate inflammatory responses driven by excessive M1 macrophage polarization, further contributing to improved patient survival.

## Supporting information

Suplementary Results

## Conflict of Interest

Thel authors declare no conflict of interest.

## 6. APPENDICE

## Acknowledgments and Sources Funding

The authors thank Prof Dr Flavia C. Meotti, Dr Larissa A. C. Carvalho and MSc Cheila J. C. Gomes, from the Institute of Chemistry – University of São Paulo, for donation and authentication of HL-60 and THP-1 cell lines, as well as some antibodies and other reagents. We thank Prof Dr Carlos F. M. Menck and the technician Dr Veridiana Munford, from the Institute of Biomedical Sciences – University of São Paulo, for supporting us with the Accuri cytometer analysis. We thank Prof Dr Alexandre B. Cardoso and Dr Lilian C. Russo, from the Institute of Chemistry – University of São Paulo, and Prof Dr Helena B. Nader, from INFAR -Paulista School of Medicine – Unifesp, for the use of the microscopes Leica DMi8 widefield and Zeiss LSM 780 confocal, respectively. We also thank Prof. Bianca S. Zingales and the technician Marcelo N. Silva, from the Institute of Chemistry - University of São Paulo, for equipment usage and reagents exchange. This work was primarily supported by grants from the Sao Paulo Research Foundation - FAPESP (Grants No 2015/03983-0, No 2018/01753-6 and 2022/04243-4) and the Brazilian National Research Council – CNPq (Grant No 402230/2016-7 and 304358/2021-5) to FLF. JOF was supported by a FAPESP PhD fellowship (Grant No 2017/16491-4).

## Author Contributions

JOF designed and performed the experimental study; JOF, DRDCGP and YTM performed bioinformatic analysis; VKB and DJFH analyzed the text and figures format; FLF designed and supervised the study; JOF, DRDCGP, YTM, VKB, DJFH and FLF analyzed and interpreted the data; JOF, DRDCGP, VKB and FLF wrote the final manuscript.

## Availability of data and material

All data raised in this work are presented in the main figures of the manuscript and in the supplementary data.

---

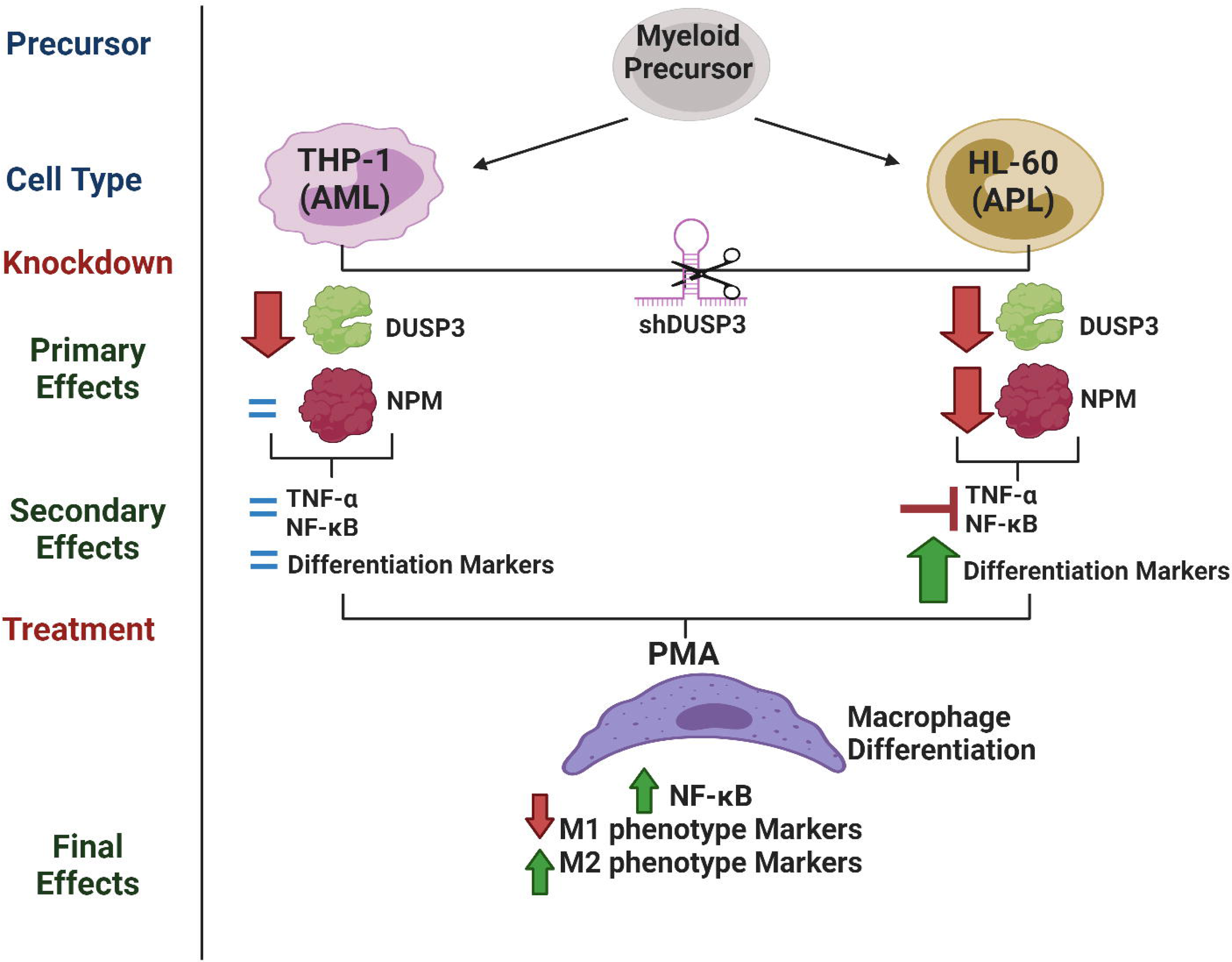

## Notes

### Competing Interest Statement

The authors have declared no competing interest.

