## Supplementary material for "Dual Specificity Phosphatase 3 knockdown drives myeloid leukemia cells to differentiate into macrophages and polarize": Suplementary Results

**Supplementary Materials**
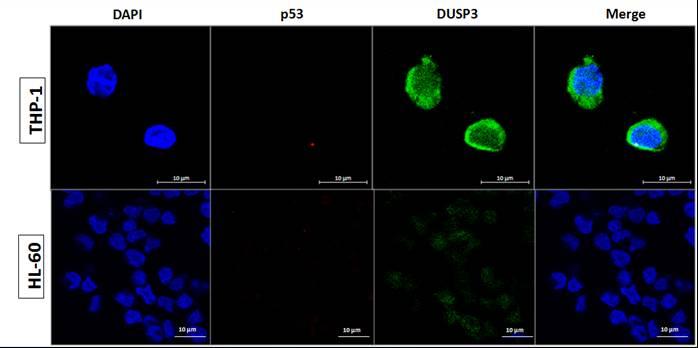

**Supplementary Figure S1. Immunofluorescence analysis of DUSP3 and p53 in THP-1 and HL-60 cells.** DUSP3 is differentially expressed in different cell lines of acute myeloid leukemia. In agreement with immunoblotting results, immunofluorescence analysis revealed that DUSP3 expression (green staining) is higher in untreated and undifferentiated THP-1 cells when compared to HL-60 cells, while p53 expression (red staining) was not detected in either cell line.

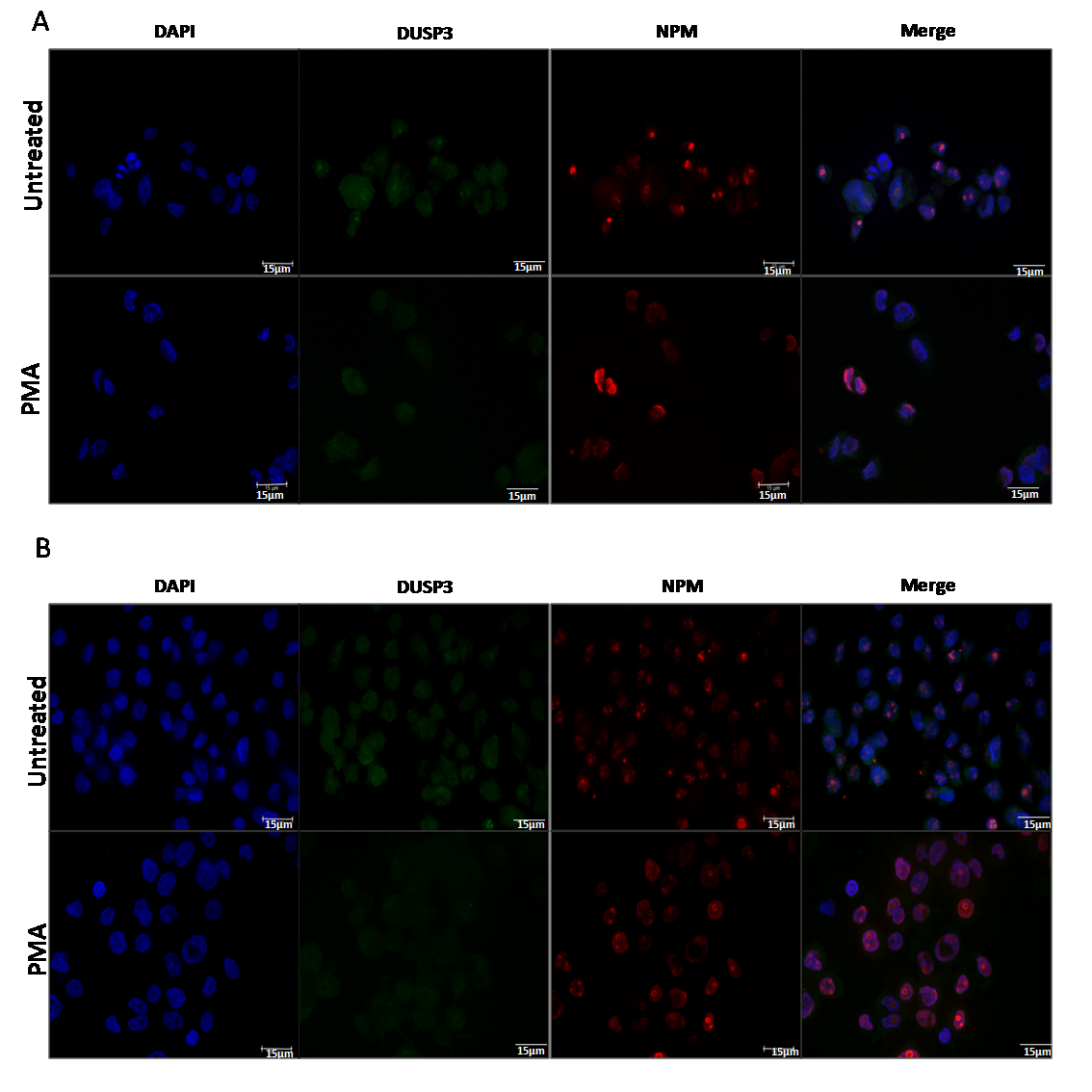

**Supplementary Figure S2. Immunofluorescence analysis of DUSP3 and NPM in THP-1 and HL-60 cells, with or without PMA-induced differentiation.** Consistent with immunoblotting results, immunofluorescence analysis showed that: (A) In THP-1 cells, both untreated (top) and PMA-treated (bottom) for macrophage differentiation, there were no variations in DUSP3 (green staining) and NPM (red staining) protein expression. (B) However, untreated HL-60 cells (top) exhibited higher expression of DUSP3 and NPM compared to those treated with PMA (bottom).

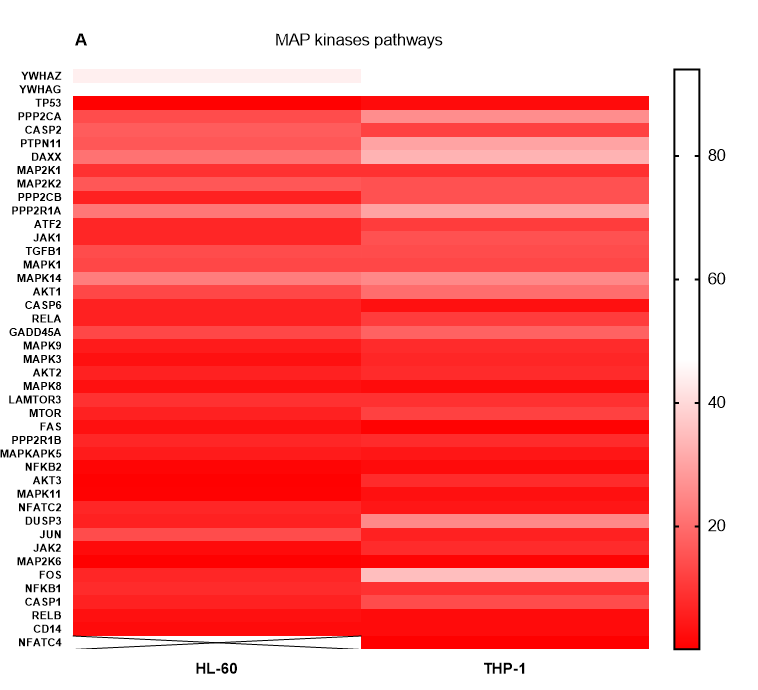

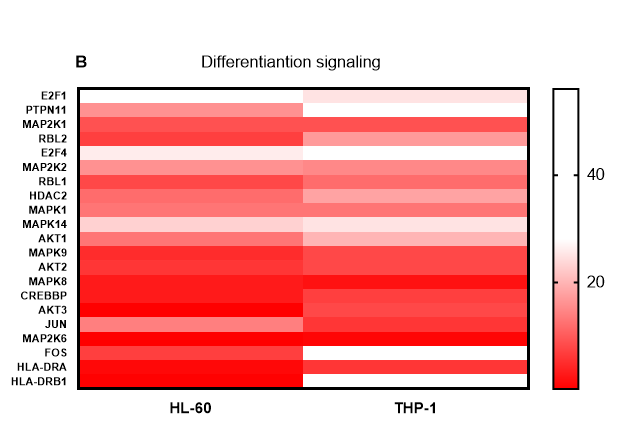

**Supplementary Figure S3. Expression of key genes involved in the MAP kinase pathway and macrophage differentiation, expressed as fragments per kilobase of transcript per million mapped reads (FPKM).** Consistent with immunoblotting results, bioinformatics analysis indicated that: (A) HL-60 cells have higher *dusp3* gene expression compared to THP-1 cells. (B) In the differentiation pathway of these cell lines, other genes, primarily phosphatases and kinases, were also differentially expressed.

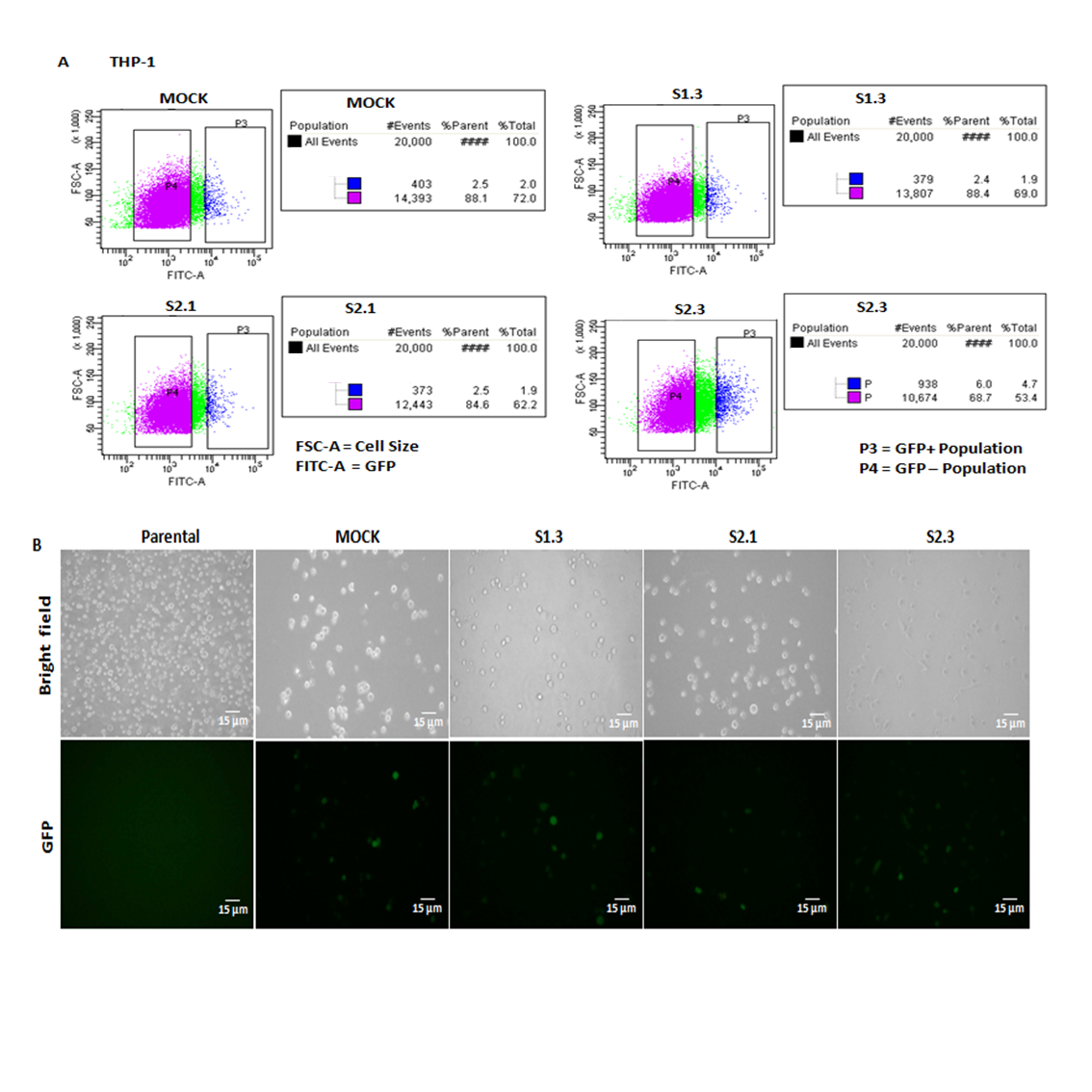

**Supplementary Figure S4. GFP fluorescence-based sorting of THP-1 cells after lentiviral transduction with shDUSP3.** (A) Transduced populations with Mock and shDUSP3 vectors were separated using the FACS-Aria III Cell Sorter via the GFP-tag in the FITC channel. (B) One day post-sorting, cells were monitored using an inverted microscope with a GFP filter in contrast to bright-field imaging.

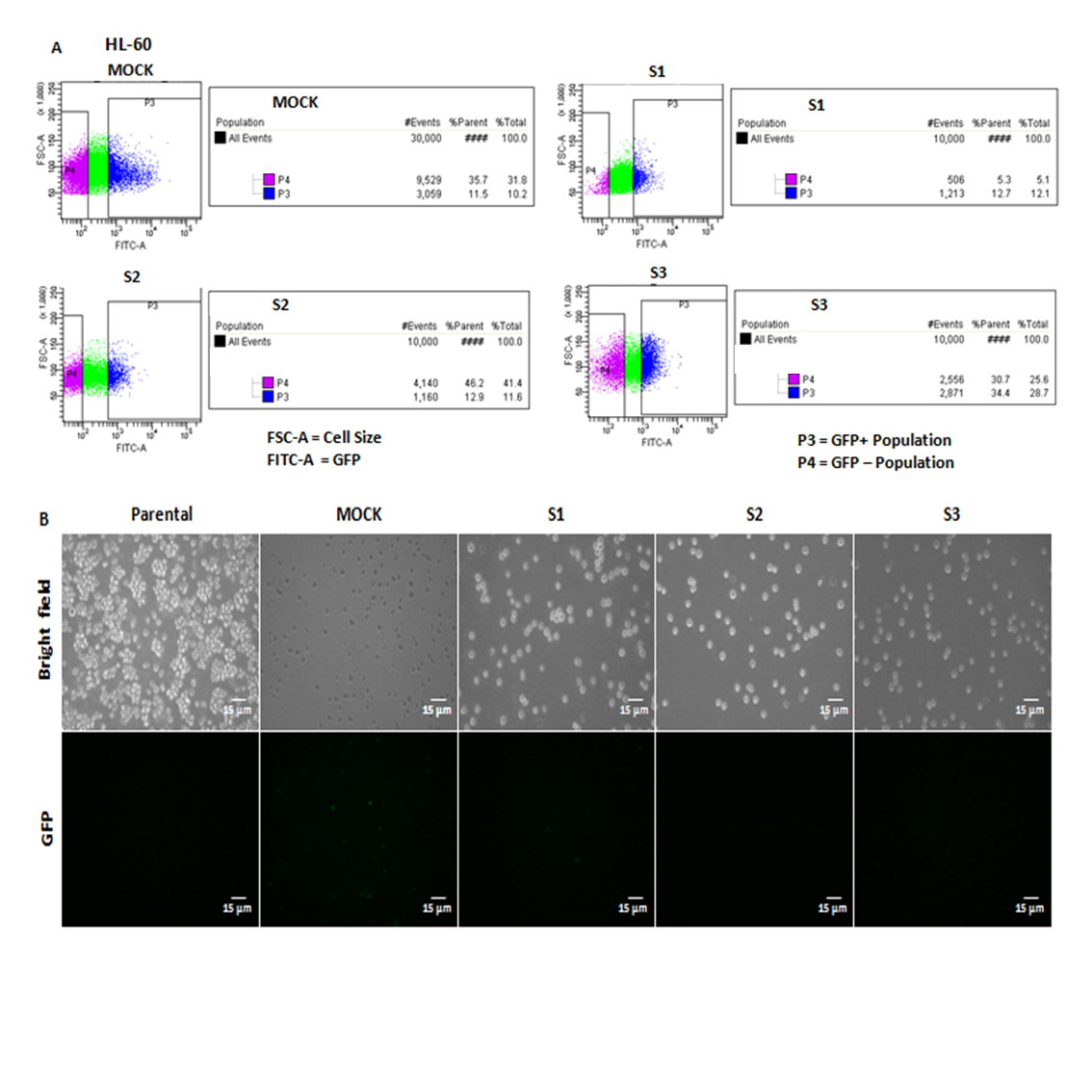

**Supplementary Figure S5. GFP fluorescence-based sorting of HL-60 cells after lentivirus transduction with shDUSP3.** (A) Transduced populations with Mock and shDUSP3 vectors were sorted using the FACS-Aria III Cell Sorter via the GFP-tag in the FITC channel. (B) One day post-sorting, cells were monitored using an inverted microscope with a GFP filter in contrast to bright-field imaging.

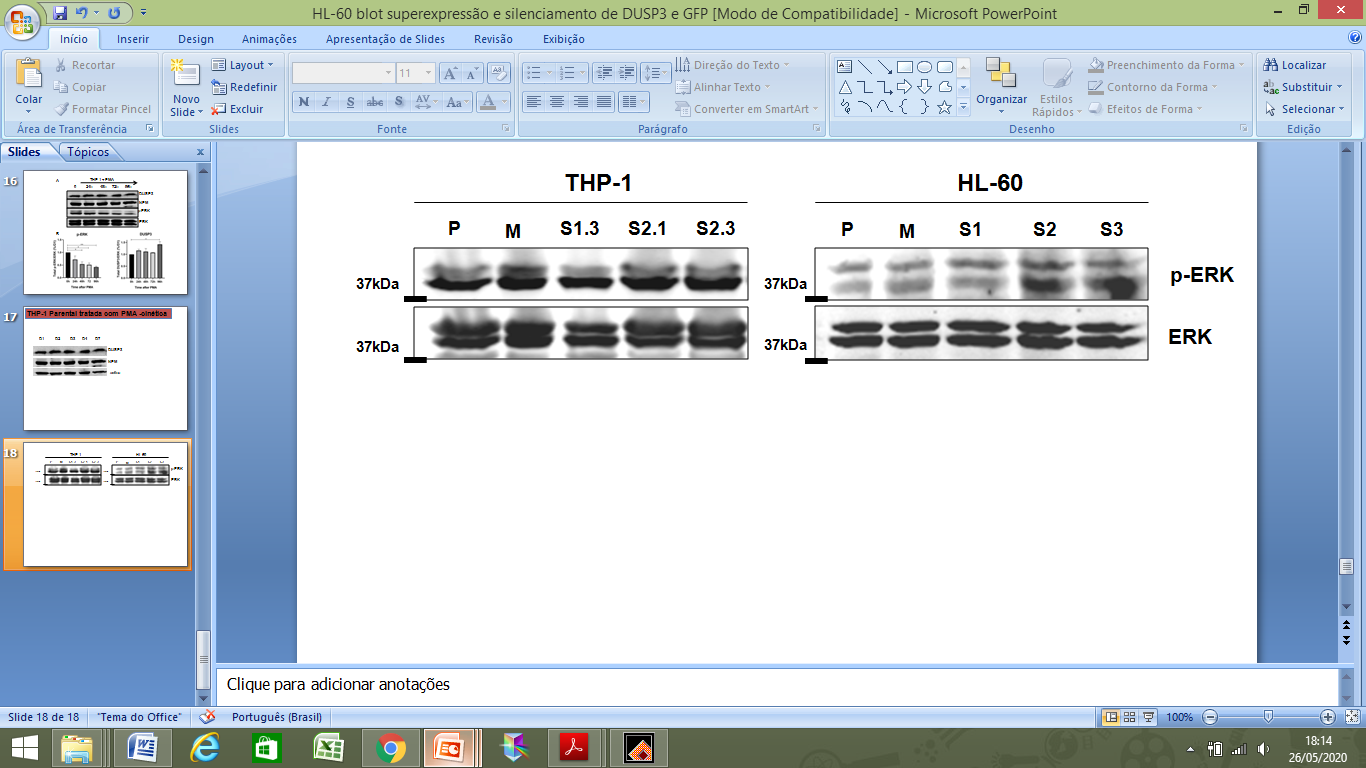

**Supplementary Figure S6. Analysis of p-ERK1/2 expression before and after DUSP3 knockdown.** Western blot analysis was performed to assess p-ERK1/2 expression in THP-1, and HL-60 cell lines before and after DUSP3 knockdown. The changes in p-ERK1/2 expression were minimal and not statistically significant in either cell line. Immunoblots representative of three independent experiments.

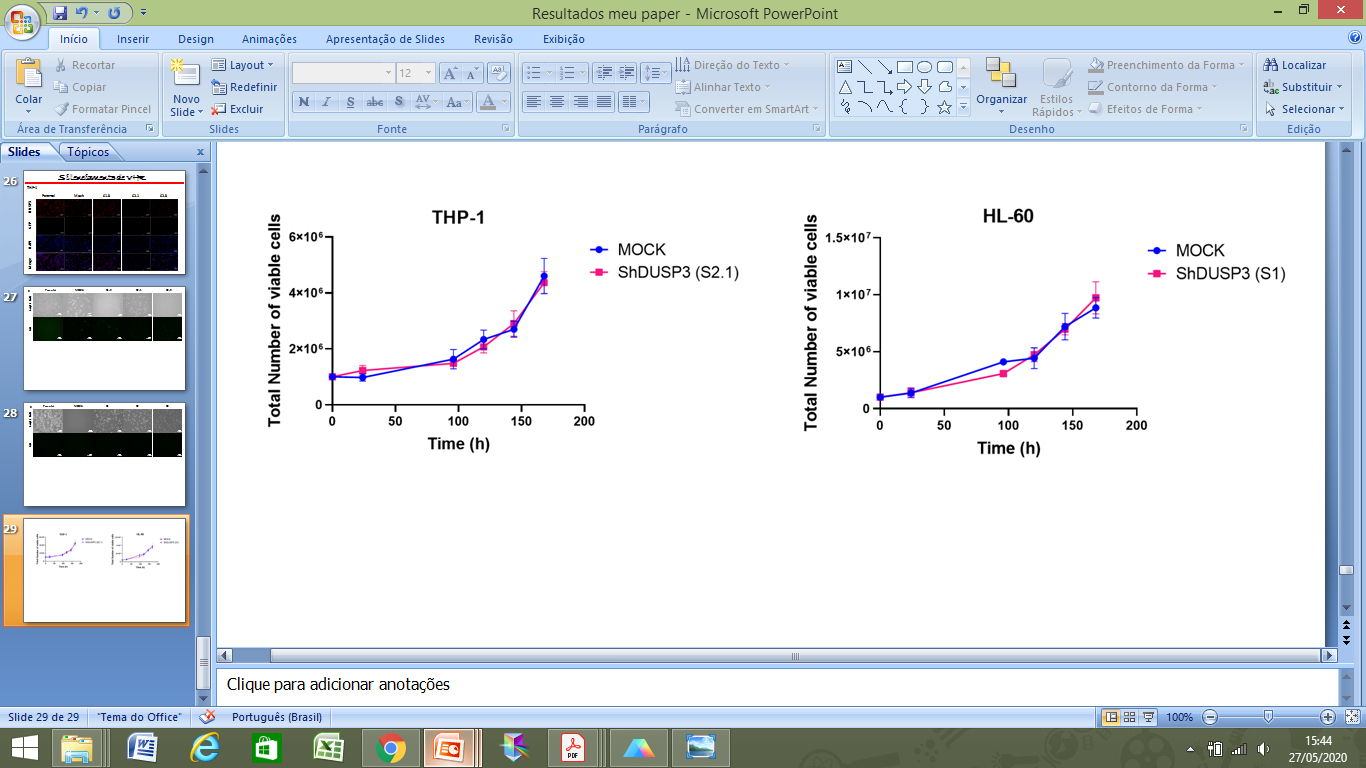

**Supplementary Figure S7. Proliferation curves of THP-1 and HL-60 cells with or without DUSP3 knockdown.** There were no differences in the proliferation of THP-1 or HL-60 cells with DUSP3 knockdown compared to non-silenced controls. Cells were plated at a density of 1x10^6 at 0h and counted at 24, 96, 120, 144, and 168h. Data from three independent experiments are presented as mean ± SD.

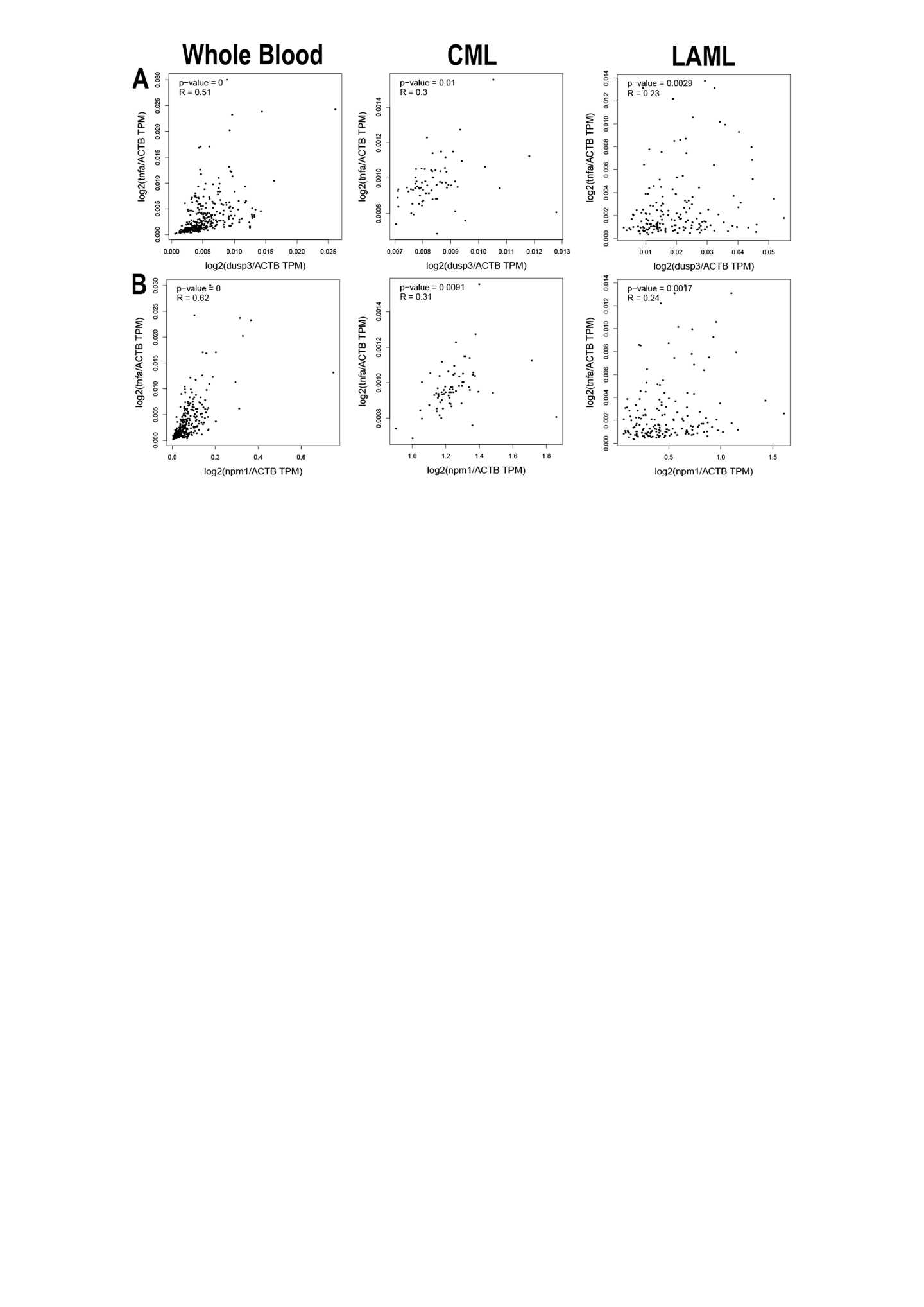

**Supplementary Figure S8. Correlation analysis of *dusp3*, *npm1*, and *tnfa* gene expression by Pearson’s correlation.** (A) In CML and AML (= LAML) samples there is a weak positive correlation between the expression of the *dusp3* and *tnfa*, which becomes negative in whole blood. (B) The correlation between *npm1* and *tnfa* expression is positive in mature blood cells and decreases in different types of leukemia (CML and LAML, respectively). Analyses were conducted using GEPIA2 with data from TCGA Cancer and GTEx.

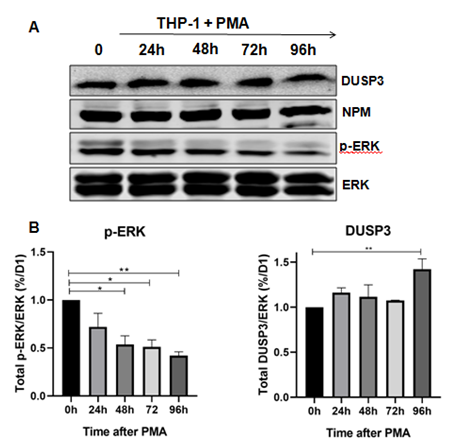

**Supplementary Figure S9. Kinetics of DUSP3, NPM and p-ERK1/2 proteins expression following PMA-induced differentiation of THP-1 cells.** (A) Parental THP-1 cells were treated with PMA to induce macrophage differentiation, and samples were lysed for Western blotting at different time points post-treatment (24h, 48h, 72h, and 96h). A slight change in DUSP3 and a decrease in p-ERK1/2 expression were observed following differentiation, while NPM expression remained unchanged. (B) Quantification of DUSP3 and p-ERK1/2 expression was performed by densitometry using Image Studio Lite Ver 5.2. Band intensities of DUSP3 and p-ERK1/2 were normalized to total ERK. Fold changes in DUSP3 and p-ERK1/2 levels are expressed relative to levels in undifferentiated cells. Data are presented as mean ± SD of three independent experiments.
